# Neuroligin dependence of social behaviour in *C. elegans* provides a model to investigate an autism associated gene

**DOI:** 10.1101/2020.02.03.931592

**Authors:** Helena Rawsthorne, Fernando Calahorro, Emily Feist, Lindy Holden-Dye, Vincent O’Connor, James Dillon

**Affiliations:** School of Biological Sciences, Highfield Campus, University of Southampton, Southampton, SO17 1BJ, UK

## Abstract

Autism spectrum disorder (ASD) is characterised by a triad of behavioural impairments including social behaviour. Neuroligin, a trans-synaptic adhesion molecule, has emerged as a penetrant genetic determinant of behavioural traits that signature the neuroatypical behaviours of autism. However, the function of neuroligin in social circuitry and the impact of genetic variation to this gene is not fully understood. Indeed, in animal studies designed to model autism there remains controversy regarding the role of neuroligin dysfunction in the expression of disrupted social behaviours. The model organism, *C. elegans,* offers an informative experimental platform to investigate the impact of genetic variants on social behaviour. In a number of paradigms it has been shown that inter-organismal communication by chemical cues regulates *C. elegans* social behaviour. We utilise this social behaviour to investigate the effect of autism associated genetic variants within the social domain of the research domain criteria. We have identified neuroligin as an important regulator of social behaviour and segregate the importance of this gene to the recognition and/or processing of social cues. We also use CRISPR/Cas9 to edit an R-C mutation that mimics a highly penetrant human mutation associated with autism. *C. elegans* carrying this mutation phenocopy the behavioural dysfunction of a *C. elegans* neuroligin null mutant, thus confirming its significance in the regulation of animal social biology. This highlights that quantitative behaviour and precision genetic intervention can be used to manipulate discrete social circuits of the worm to provide further insight to complex social behaviour.

## Introduction

Autism spectrum disorder (ASD) is a pervasive developmental disorder (1). It is clinically characterised by a triad of neuroatypical behaviours that include impaired verbal communication, repetitive behaviours and impaired social interactions. Sensory processing deficits span across the autism spectrum (2). For example, recognition of social cues and multi-sensory integration of those cues are often impaired (2, 3). There is a strong genetic association in ASD with hundreds of genes implicated in the disorder (4). Many of these genes encode synaptic proteins (5–7), suggesting that synaptic dysfunction underpins the expression of ASD associated neuroatypical behaviours (5). However, it is still unclear how genetic variants lead to changes in neural circuits that result in the spectrum of characteristics that are represented in individuals with a recognizable ASD diagnosis. The role of individual genes and the wider complex interaction between polygenic loci make the genetic architecture of ASD complex. This complexity requires investigation that will ultimately delineate the weight of contribution in genetic backgrounds that range from a single penetrant gene to ones with multiple common variants at multiple loci (8).

One of the synaptic proteins that have been implicated in ASD is neuroligin (NLGN). NLGN is a post-synaptic cell adhesion protein which aids the stabilisation of synaptic function (9). One highly penetrant variation which has been shown to affect the NLGN3 gene of autistic individuals is the R451C mutation (10). This missense variant causes a substitution of arginine for cysteine in the extracellular domain of NLGN3 which leads to its misfolding and failure to traffic to the cell surface. It has been suggested that the R451C mutation may result in a loss of function and disruption of synaptic function and plasticity within a number of central neural circuits (11). Despite broad investigation it remains unclear as to how penetrant this mutation and further neuroligin mutants are in disrupting social behaviour in animal models. There have been differing reports of disrupted social behaviour in mice where the mutation has been introduced into strains with different genetic backgrounds (12, 13).

The model organism, *Caenorhabditis elegans,* is a good experimental platform to investigate the effect of ASD associated genetic variants. *C. elegans* facilitate systems level analysis due to their genetic tractability, simple nervous system and gene homology to humans (14, 15). Included within this is the conservation of genes involved in synapse maturation and function (16–18). *C. elegans* encode a single orthologue of mammalian NLGN3 called *nlg-1*, which has been shown to share key structural and functional domains with the human protein (19, 20). This conservation is reinforced by observations showing human NLGN is able to provide functional rescue of a *nlg-1* deficiency in *C. elegans* (19).

In order to investigate the potential function of *nlg-1* within the social domain, we have utilised a paradigm of inter-organismal communication between *C. elegans* and progeny. The effect of chemosensory stimuli on worm behaviour can be investigated using an assay in which the propensity of an adult worm to leave its food source, a bacterial lawn, is monitored over time. This food leaving assay scores food leaving events in which each event is defined as an occasion when the whole of the worm’s body comes off food. This is a behavioural output which can be modulated in response to different cues (21). For example, when exposed to ad libitum source of food, *C. elegans* will remain on the food lawn and perform infrequent food leaving events (21). However, an increase in the number of progeny populating an otherwise replete food lawn causes a population-dependent increase in food leaving events of adult worms. This progeny induced leaving is not observed in a *daf-22* loss of function mutant (22) showing that offspring produced ascaroside pheromones modulate adult behaviour. This suggests that inter-organismal signalling from progeny to parents on food replete lawns provides a quantifiable behaviour to investigate the genetic determinants of social circuits (22).

We use this assay to show that the ASD associated gene, NLGN, is an important determinant in regulating *C. elegans* social circuitry. We identify that a genetic change that models the human R451C phenocopies the functional null. In doing this, we facilitate further insight into the genetic underpinnings of social behaviour, a key diagnostic Research Domain Criteria in autism and other psychiatric disorders (https://www.nimh.nih.gov/research/research-funded-by-nimh/rdoc/index.shtml).

## Results

### The importance of *nlg-1* in modulating social communication in adult worms

Two of the major characteristics of ASD are impaired social behaviour and deficits to multi-sensory integration (2). It has been previously shown that *C. elegans* elicit a social response to progeny populating a food lawn (22). More specifically, in response to increasing numbers of progeny on food, wild-type (N2) worms show a population-dependent increase in food leaving in the presence of an otherwise replete food lawn. This progeny enhanced food leaving behaviour was shown to be the result of chemosensory social signalling between progeny and adult worms (22). We have used this social behaviour to investigate the autism associated gene, NLGN3. To investigate the behaviour of *nlg-1(ok259)* null mutants in response to progeny, worms were picked onto the centre of a bacterial lawn and food leaving events were counted at 2 and 24 hours. At 2 hours no progeny are present on the food lawn. In comparison, at 24 hours progeny will have accumulated on the food lawn. Therefore, quantifying food leaving events at these time points allows for a direct comparison of food leaving behaviour in the absence and presence of progeny.

Observing food leaving in N2 adults showed an increase in leaving events over time. After 2 hours, when no progeny were present, worms remained on the food lawn (Figure 1A). At 24 hours, when the food lawn is populated with progeny, N2 display progeny enhanced food leaving behaviour, in which they show a 12-fold increase in the number of food leaving events (Figure 1A). *nlg-1(ok259)* mutants remained on food, similar to N2, after 2 hours on the food lawn (Figure 1A). However, at 24 hours, when the food lawn is dense with progeny, *nlg-1(ok259)* leaves the food patch less than N2 (Figure 1A). *nlg-1(ok259)* did not show the same degree of progeny enhanced food leaving as N2, suggesting that their social interaction with progeny may be impaired.

**Figure 1:**
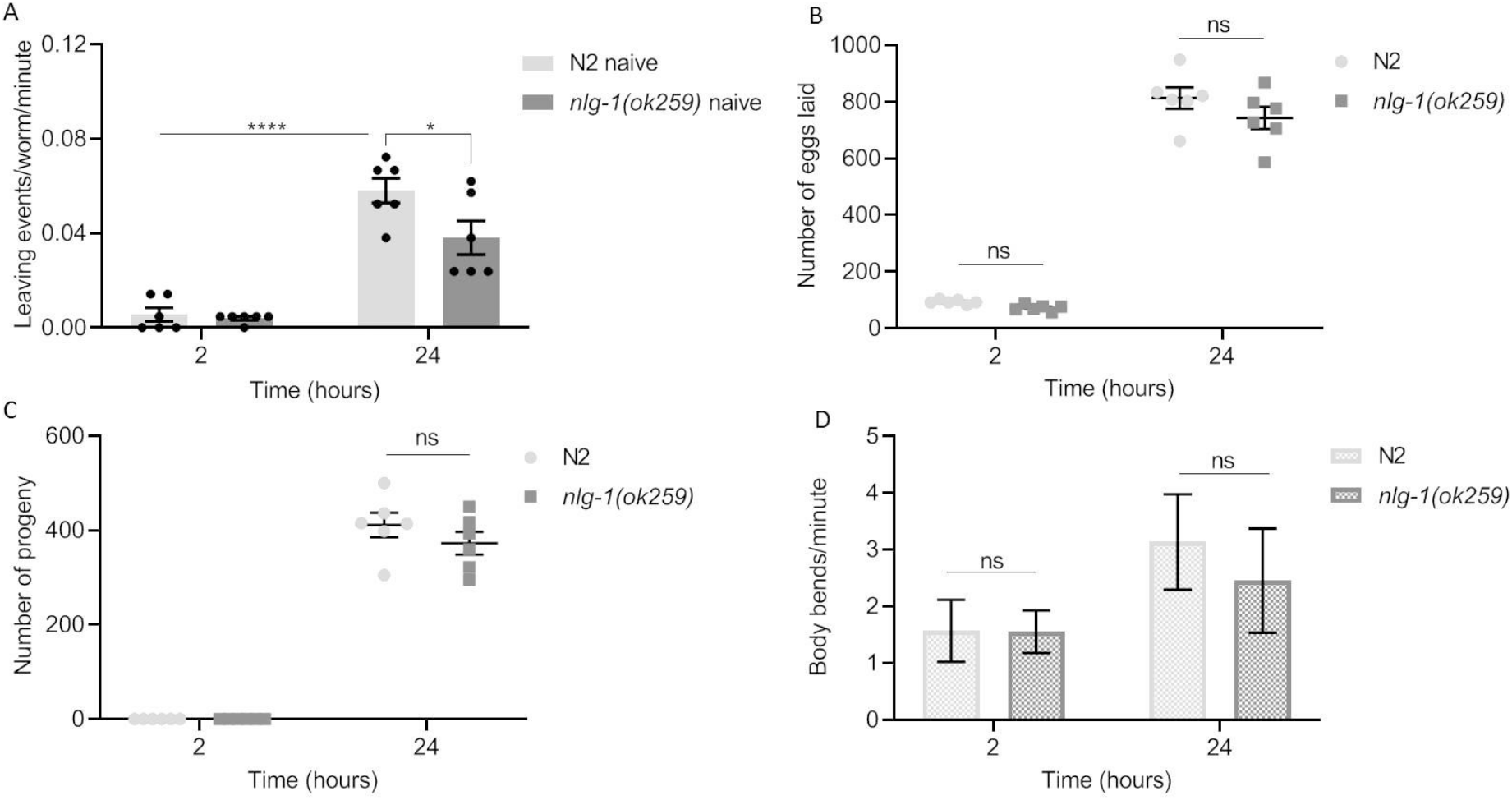
*nlg-1(ok259)* adults show deficient food leaving in response to progeny. (A) A food leaving assay was performed with N2 and *nlg-1(ok259)* null mutant. N2 and *nlg-1(ok259)* adults were picked onto the centre of a bacterial lawn before food leaving events were counted at 2 and 24 hours. N2 and *nlg-1(ok259)* n=6. (B and C) the number of eggs (B) and progeny (C) for N2 and *nlg-1(ok259)*. N2 and *nlg-1(ok259)* n=6. (D) The number of body bends per minute performed by N2 and *nlg-1(ok259)* worms on food at 2 and 24-hours. N2 n=6. *nlg-1(ok259)* n=4. All data shown as mean ±SEM. Statistical analysis performed using Two-way ANOVA with Sidak’s multiple comparison test; ns P≥0.05, * P<0.05, **** P≤0.0001.

We next wanted to understand if the reduced food leaving behaviour of *nlg-1(ok259)* was due to impaired social interaction with progeny. We reasoned that reduced food leaving could also result if *nlg-1(ok259)* had an egg laying deficiency. Previous studies had shown that the amount of food leaving was directly related to the number of progeny, with a reduced progeny accumulation giving a reduced food leaving (22). To test this, the number of eggs laid and the number of progeny present on the food lawn was counted after each food leaving assay. For both N2 and *nlg-1(ok259)* the total number of eggs laid (Figure 1B) during the assay, and number of progeny present at 24-hours were counted (Figure 1C) and there was no difference between the strains. These data show the reduced food leaving of *nlg-1(ok259)* cannot be explained by reduced progeny available to drive adult food leaving. Furthering this, no difference was seen in the number of body bends for *nlg-1(ok259)* (Figure 1D) indicating that locomotion per se does not underlie the reduced food leaving of *nlg-1(ok259)*. Thus, the deficit in progeny enhanced food leaving behaviour of *nlg-1(ok259)* appears due to impaired social interaction of the mutant with progeny.

Next, we wanted to investigate whether the reduced food leaving described above for *nlg-1(ok259)* was due to an impaired social interaction with progeny. To this end, we used a pre-conditioned food lawn, as previously described (22). Pre-loading progeny onto a food lawn before assaying the adult food leaving events preconditions the lawn with chemosensory cues released by progeny. This recapitulates the progeny dense conditions worms are exposed to after 24-hours on the food lawn but the assayed adults undergo more acute exposure to the progeny (Figure 2A).

**Figure 2:**
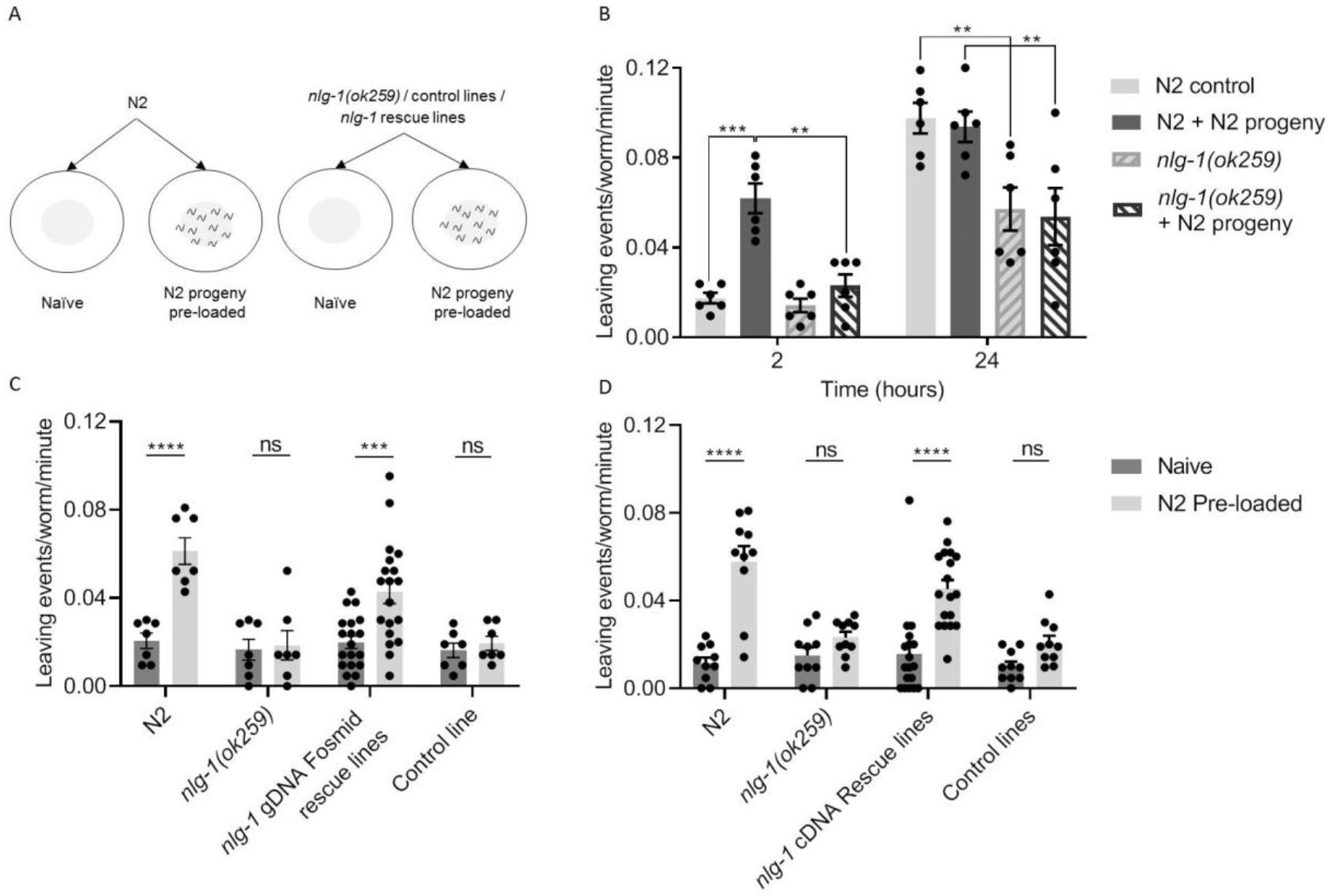
*nlg-1(ok259)* show a deficit in social interaction when exposed to N2 progeny. (A) Cartoon showing the principles of the pre-conditioned food leaving assay. In order to test the effect of progeny on food leaving behaviour, *C. elegans* adults were picked onto either a naïve or pre-conditioned food lawn. A naïve lawn contains no progeny whereas a pre-conditioned food lawn contains ~140 N2 progeny. (B) N2 and *nlg-1(ok259)* were picked onto naïve and pre-conditioned food lawns and their food leaving behaviour observed at 2 and 24 hours. N2 and *nlg-1(ok259)* n=6. Two-way ANOVA with Tukey’s multiple comparison test; ** P≤0.01, *** P≤0.001. (C) The number of food leaving events of N2, *nlg-1(ok259)*, *nlg-1* gDNA fosmid rescue *nlg-1(ok259) X, Ex* [WRM0610dD09; P*myo-3*∷*gfp*] and *nlg-1(ok259) X, Ex* [pPD95.77; P*myo-3∷gfp*] control adults on naïve and pre-conditioned food lawns. N2, *nlg-1(ok259)* and *nlg-1(ok259) X, Ex* [pPD95.77; P*myo-3∷gfp*] control line n=7, *nlg-1(ok259) X, Ex* [WRM0610dD09; P*myo-3*∷*gfp*] n=19. Two-way ANOVA with Sidak’s multiple comparison test; ns P≥0.05, *** P≤0.001, **** P≤0.0001. (D) The number of food leaving events of N2, *nlg-1(ok259)*, *nlg-1* cDNA rescue *nlg-1(ok259) X, Ex* [pPD95.77 (P*nlg-1∷nlg-1* Δ#14); P*myo∷gfp*] and *nlg-1(ok259) X, Ex* [pPD95.77; P*myo-3∷gfp*] control line on naïve and pre-conditioned food lawns. N2, *nlg-1(ok259)* and *nlg-1(ok259) X, Ex* [pPD95.77; P*myo-3∷gfp*] control line n=10. *nlg-1(ok259) X, Ex* [pPD95.77 (P*nlg-1∷nlg-1* Δ#14); P*myo∷gfp*] n=18. Two-way ANOVA with Tukey’s multiple comparison test; ns P≥0.05, **** P≤0.0001. All data shown as mean ±SEM

The pre-conditioning with N2 progeny results in enhanced food leaving in N2 adults compared to the naive control at 2 hours (Figure 2B). This is consistent with previous findings which showed that social interaction of adult worms and progeny on pre-conditioned food lawns results in enhanced food leaving in N2 *C. elegans* (22). In comparison, *nlg-1(ok259)* adults when exposed to plates preconditioned with N2 progeny did not show enhanced food leaving. They left infrequently on both naïve and pre-conditioned food lawns at 2-hours (Figure 2B). This confirms that *nlg-1(ok259)* have reduced food leaving in response to progeny compared to N2. Therefore, this suggests that NLG-1 may be an important regulator of chemosensory driven social interaction in *C. elegans*. To confirm the importance of NLG-1 in the social circuit of the worm we generated two transgenic rescue lines expressing either *nlg-1* gDNA or *nlg-1* cDNA in the *nlg-1(ok259)* background. In response to pre-loaded N2 progeny both the rescue lines expressing *nlg-1* gDNA and cDNA showed progeny enhanced food leaving, similar to that of N2 (Figure 2C and D). This shows that expression of either *nlg-1* in its genomic or cDNA form can rescue the reduced food leaving of *nlg-1(ok259)* mutants in response to progeny. Together, these data suggest that *nlg-1(ok259)* mutants are impaired in their ability to modulate food leaving behaviour in the presence of progeny and that NLG-1 may play an important role in regulating this social behaviour.

We next wanted to segregate whether NLG-1 is important for the production and/or release of social cues from progeny or the recognition and/or integration of the social cue in the adult worm. To do this we pre-conditioned food lawns with either N2 or *nlg-1(ok259)* progeny. Food leaving behaviour was then observed for N2 and *nlg-1(ok259)* adults in response to either N2 or *nlg-1* progeny (Figure 3A). In this way we were able to investigate whether *nlg-1(ok259)* mutant progeny are capable of driving enhanced food leaving behaviour in adult worms, hence informing on their ability to produce / release chemical social cues.

**Figure 3:**
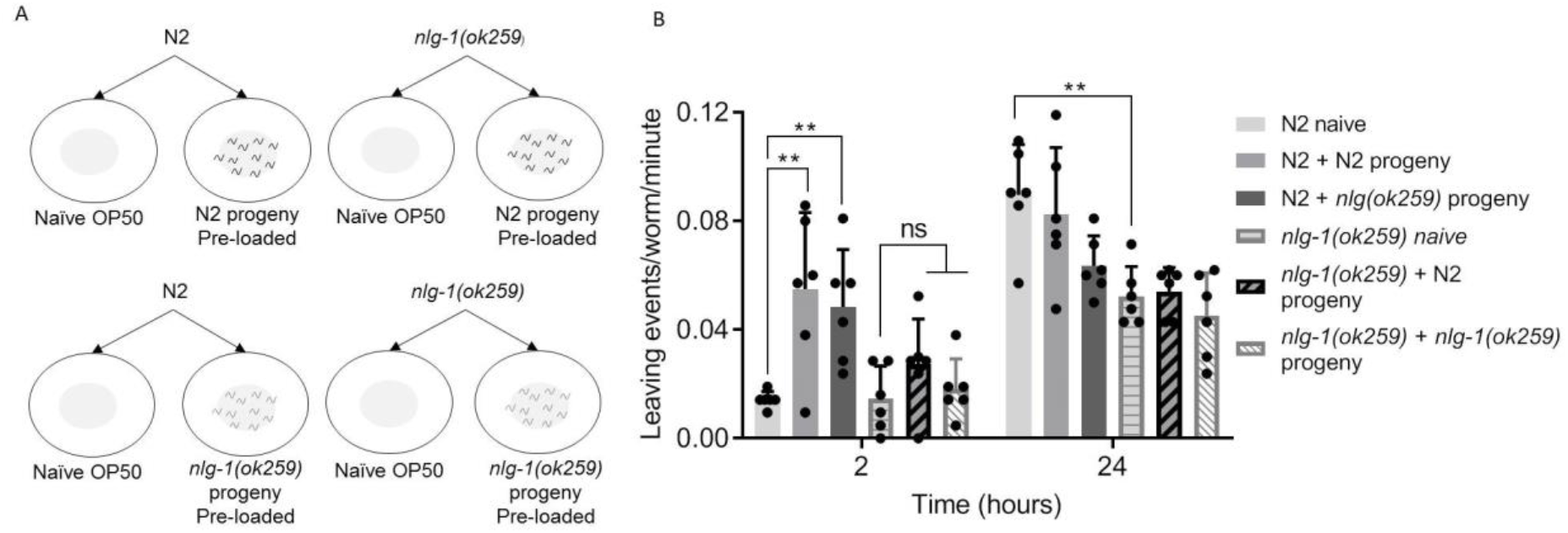
Parental *nlg-1* is required for recognition and/or processing of progeny derived social cues. (A) Experimental setup. N2 and *nlg-1(ok259)* were placed onto a naïve OP50 bacterial lawn or a lawn pre-conditioned with N2 or *nlg-1(ok259)* progeny. (B) The number of food leaving events of N2 and *nlg-1(ok259)* adults on naïve and pre-conditioned food lawns. Data are mean ±SEM. N2 and *nlg-1(ok259)* n=6. Two-way ANOVA with Tukey’s multiple comparison test; ns P≥0.05, ** P≤0.01.

N2 adults showed enhanced food leaving behaviour in the presence of both N2 and *nlg-1(ok259)* progeny (Figure 3B). This suggests that both N2 and *nlg-1(ok259)* progeny are capable of driving enhanced food leaving behaviour in N2 adults. In turn, this suggests that *nlg-1(ok259)* mutant progeny are capable of producing and releasing chemical social cues in order to stimulate enhanced food leaving in adults. In comparison, *nlg-1(ok259)* adults do not show enhanced food leaving in response to either progeny relative to the naïve control at 2-hours (Figure 3B). Taken together, these results suggest that N2 and *nlg-1(ok259)* progeny are not impaired in the production and release of social cues. Furthermore, these results are consistent with the hypothesis that *nlg-1(ok259)* adults have impaired social communication with progeny which may involve deficits in recognition and/or integration of progeny derived social cues.

### *C. elegans* carrying R433C mutation phenocopy the social impairment of *nlg-1* null

So far we have shown that the *nlg-1(ok259)* null mutant does not show progeny driven enhanced food leaving behaviour and we hypothesise that this behavioural deficit is specific to the recognition and/or integration of social cues in the adult worm. In this way, *nlg-1(ok259)* mutants are modelling neuroatypical behaviour in the social domain. Considering this, we next wanted to know whether *C. elegans* could model the same social impairment but in response to a specific human genetic variation implicated in autism. We investigated a penetrant missense variant which has been identified in the NLGN3 gene of individuals on the Autistic spectrum. The R451C mutation results in an R-C amino acid substitution within the extracellular cholinesterase-like domain of NLGN3 (10). *C. elegans* encode a conserved arginine within the cholinesterase-like domain of NLG-1 which is present in human NLGN1-4 (Figure 4A). Using CRISPR/Cas9 the R451C mutation was generated in *C. elegans* by editing an R-C substitution at position 433 within the cholinesterase-like domain of NLG-1 (Figure 4A). The bona fide nature of this CRISPR event was confirmed by genomic sequencing which identified that the CGA codon of the wild type was converted to TGC encoding cysteine in the mutant line (Supplementary Figure 1). The conservation in sequence suggests that the same disruption in NLGN structure that arises in the human protein with this mutation at position 451 will be replicated in *C. elegans* with the orthologues mutation at 433 (based on NLG-1 isoform C40C9.5e). The social behaviour of the CRISPR generated line, *nlg-1(qa3780),* was investigated using lawns that were pre-conditioned with N2 progeny in order to compare the response of CRISPR generated *nlg-1(qa3780)* to *nlg-1(ok259)* null mutant in response to N2 derived social cues.

**Figure 4:**
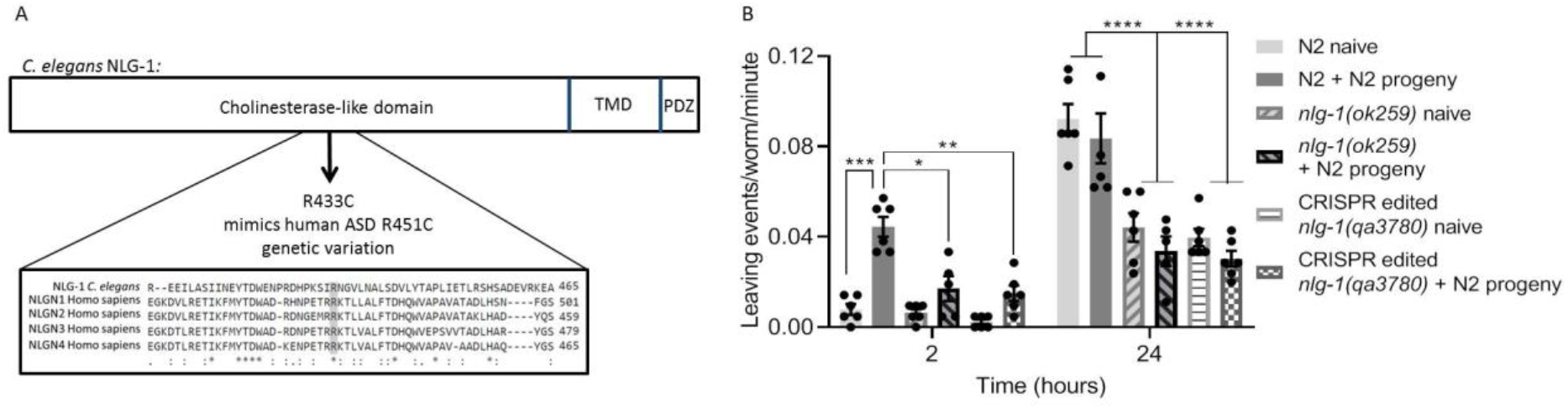
R433C mutation in *nlg-1* phenocopies social impairment of *nlg-1* null. (A) Domain structure of NLG-1 indicating the arginine to cysteine (R-C) amino acid substitution at position 433 generated within the cholinesterase like domain following CRISPR/Cas9. Protein sequence alignment of *C. elegans* NLG-1 and human NLGN1-4 indicates the arginine residue involved in ASD and its conservation in *C. elegans* (shaded in grey). ‘*’ indicates conservation of a single amino acid residue, ‘:’ indicates conservation between amino acid groups with similar properties and ‘.’ indicates conservation between amino acid groups with weakly similar properties. (B) The number of food leaving events were counted for N2, *nlg-1(ok259)* and CRISPR generated *nlg-1(qa3780)* on naïve and N2 progeny conditioned food lawns. Data are mean ±SEM. N2, *nlg-1(ok259)* and *nlg-1(qa3780)* n=6. Two-way ANOVA with Tukey’s multiple comparison test; * P<0.05, ** P≤0.01, *** P≤0.001, **** P≤0.0001.

Consistent with our previous findings, N2 adults show enhanced food leaving in response to progeny on pre-conditioned food lawns (Figure 4B). Furthermore, *nlg-1(ok259)* do not show enhanced food leaving in the presence of progeny (Figure 4B). The response of *nlg-1(qa3780)* mutants, carrying the R433C mutation, is very similar to that of *nlg-1(ok259)* null mutants. *nlg-1(qa3780)* showed no enhanced food leaving when exposed to progeny (Figure 4B). This suggests that the single R433C missense variant to *nlg-1* results in social impairment which phenocopies that of the *nlg-1* null mutant. Furthering this, these results suggest that *C. elegans* can model social impairment in response to autism associated human genetic variants.

## Discussion

ASD causes neuroatypical social behaviour and deficits in sensory processing (2). ASD has a well characterized genetic dependence, with the underlying determinats becoming increasingly well resolved with the use of quantitative genetics. There is a range of determinants that imply polygenic interaction between common variation of the genome. In addition, there are a number of more penetrant single gene mutations that are identified as contributing to the expression of behavioral traits that siganature neuroatypical behaviour of ASD. Many of these penetrant genes encode synaptic proteins (4) highlighting synaptic dysfunction as orchestrating the phenotypes associated with ASD. In the case of penetrant genes, animal experiments utilizing functional nulls or engineered mutants designed to mimic human mutations have facilitated investiagtion of the cellular circuit and system level mechanisms that disrupt behavioral domains that model ASD associated behaviour (23). In this study we have shown that *C. elegans* can be used to model social impairment in response to genetic models of the human variants identified in individuals with an ASD diagnosis. This is highlighted by our investigation and comparison of the single neuroligin gene in *C. elegans* that has high conservation with the 5 human neuroligin genes, particulalry the NLGN3 gene that has been strongly implicated in ASD (19, 24).

NLGNs are a family of synaptic adhesion proteins required for synaptic maturation through interaction with synaptic adhesion partners. Of these, the most studied interaction is between neuroligin and neurexin which can co-ordinate the localization of receptors at inhbitory synapses (25). It has been shown in *C. elegans* that this interaction and its function in the organisation of receptors is conserved at the neuromuscular junction of the worm (26). The NLGN functional null used in our study has been previously reported as having defects in synaptic transmission at the neuromuscular junction, in the absence of an obvious locomotory phenotype (26). We reached a similar finding, that the NLGN null mutation did not impair locomotion. Hence, we propose that the difference in food leaving behaviour of the null cannot be explained by impaired motor control but are mindful that there may be differences in the relative profile of more subtle locomotory sub-behaviours that underpin the food leaving behaviour in response to social cues.

The clinical data for the NLGN3 R451C mutation suggests it causes severe autism and Aspergers’ syndrome in a sibling carrying the mutation (10). This motivated animal experiments in which the functional null and R451C mutaion were compared. These mutants exhibit cellular changes selective for inhibitory synapses that disrupt plasticity and cause changes in learned behaviours associated with social interaction and motor control (12, 13, 27). In the former case there is a controversy as to how penetrant the mutations that mimic the human mutation are in causing a mutation dependent change in social interaction. The differential conclusions derived from mice strains raised in distinct genetic backgrounds suggests that background modifiers can affect the phenotypic output of the genetic mutation being investigated (12, 13). In this study we used a neuroligin null mutant and a CRISPR/Cas9 line in the same wild-type N2 genetic background to compare the effects of the mutation on social behaviour.

We utilized a previously characterized assay of progeny enhanced food leaving (22). Our results show that *nlg-1(ok259)* adults have impaired social communication with progeny. This deficit shows NLGN’s synaptic organising function may play a key role in the neural circuits that underlie social interaction. We provide evidence that neuroligin-dependent signalling is required in the adult for their ability to respond to the chemical cues generated by the progeny. This assay thus provides a quantitaive measure for neuroligin function in the adult. We used this to probe the integrity of the circuits driving this output utilizing a CRISPR/Cas9 mutant that models the penetrant human ASD mutation R451C. Our data show that this mutation phenocopies the deficit in food leaving seen in the null mutation which is consistent with the widely held view that this mutation is a loss of function (28). Taken together our data imply that *nlg-1* plays an important role in the circuits that organise social interaction in *C. elegans*. This would reinforce a conserved role for this class of adhesion molecule in organizing circuits that underpin social behaviour. Previous work has identified *nlg-1* function in the animals response to a number of enviromental cues including food, chemicals and temperature. Interestingly, disruption at the input level of sensory cues is an important emerging theme in human autism and future work will be required to address if the sensitivity to pheromones per se is the major determinant of the observed disruption. Equally, work in *C. elegans* has implicated changes in the balance of excitation and inhibition down stream of sensory inputs generating behavioral disruption in *C. elegans* so the results here may arise through an essential role for neuroligin at several levels of the social circuit responsible for the adult response to progeny induced food leaving (29, 30).

Previous studies have used *C. elegans* morphology and locomotion as readouts of disrupted behaviour in response to mutation to ASD associated genes (31, 32). We have extended this analysis to inform on the effect of a neuroligin variant on a phenotype that relates to one of the triad of impairments that make up the diagnostic criteria of ASD. The social interaction assay used in this study allows for quantification of ASD associated variants and their effect on social behaviour. The gene homology of *C. elegans* with mammalian systems, conservation of synaptic architecture and requirement for sensory-motor integration in the face of environmental cues provides a tractable paradigm for these investigations. Overall, the assay of social interaction in *C. elegans* has the potential to provide crucial insight into the neural circuits that underpin social behaviour and how genetic variants impact on those circuitries. This suggests it provides a robust platform to screen other genes implicated in autism and complex psychiatric disorders that exhibit underpinning disruption of the social domain.

## Materials and Methods

### *C. elegans* culturing and strains used

All *C. elegans* strains were maintained using standard conditions (33). *C. elegans* were age synchronised by picking L4 + 1 day old hermaphrodites onto a new plate 18 hours prior to the behavioural assay. Strains used: Bristol N2; VC228 *nlg-1(ok259) X* (x6 outcrossed); *nlg-1(ok259) X, Ex* [WRM0610dD09; P*myo-3*∷*gfp*]; *nlg-1(ok259) X, Ex* [pPD95.77 (P*nlg-1∷nlg-1* Δ#14); P*myo-3∷gfp*]; *nlg-1(ok259) X, Ex* [pPD95.77; P*myo-3∷gfp*]; CRISPR/Cas9 edited XA3780 *nlg-1(qa3780)* (×2 outcrossed). For transgenic animals, stable lines were selected for behavioural analysis.

### Rescue construct and transgenic methods

The *nlg-1* fosmid, WRM0610dD09, was provided by SourceBioScience. The *nlg-1* cDNA rescue construct was designed as previously described (34). Briefly, 2.5kb P*nlg-1* was cloned into the pPD95.77 vector. Subsequently, *nlg-1* Δ#14 cDNA sequence was fused to P*nlg-1* (35).

*nlg-1(ok259)* L4+1 day old worms were microinjected with *nlg-1* fosmid WRM0610dD09 (0.3ng/μl) or *nlg-1* cDNA rescue plasmid (50ng/μl) and the marker plasmid P*myo-3∷gfp* (30ng/μl). Control lines were microinjected with marker plasmid P*myo-3∷gfp* (30ng/μl).

### Behavioural assays

#### Food leaving

5cm NGM plates were prepared using a standard protocol (33). Plates were seeded with OP50 *E.coli* as described previously (22). 50μl of OP50 *E. coli* at OD_600_ of 0.8 was gently spotted on the middle of an unseeded plate the day prior to the assay. For a naive food leaving assay, seven age synchronised L4+1 day old worms were gently picked onto the centre of the bacterial lawn on the assay plate. To pre-condition the food lawn, progeny were loaded onto the bacterial lawn as previously described (22). 10 gravid adults were picked onto the bacterial lawn and left to lay 140-200 eggs before being picked off. Approximately 18-hours following this, seven L4+1 day old worms were picked onto the centre of the bacterial lawn. In all food leaving assays the number of food leaving events were counted during 30-minute observations at 2-hours only or 2 and 24-hours. A food leaving event is defined as when the whole of the worm’s body comes off the bacterial lawn. For all assays N2 animals were systematically observed in parallel with the strain under investigation. For all behavioural analysis investigators were blind to the genotypes being observed.

### Body bends

Following a food leaving assay 5-7 worms were selected for body bend quantification. Each worm was observed for 1 minute on food. A body bend was defined as a muscle contraction that resulted in a dorsal or ventral bend of the worm’s body.

### Genome editing

CRISPR/Cas9 editing was generated using a previously described method (36). N2 L4+1 day old hermaphrodites were microinjected with expression vectors for Cas9 and sgRNA’s targeting *nlg-1*, *unc-58* and *dpy-10*. The sequence of the *nlg-1* sgRNA, synthesised by Integrated DNA Technologies (IDT) was: 5’-GATTTCGAATTGATTTCGGGTGG-3’. The sequence for *unc-58* and *dpy-10* sgRNA were as previously described (37). The repair templates used for co-CRISPR genes *unc-58* and *dpy-10* were as previously described (36). The repair template targeted to *nlg-1* was: 5’-ACGCCTCAAAAATTAAAGGTAGAACATTTATTTCATCATTATAGGACCACCCGAAATCAATT**<u>TGC</u>**AAT GGAGTTCTGAATGCTCTTAGCGACGTACTTTACACCGCACCTCTCATTGAAACATTGCGAAG-3’. The mutated codon is underlined. Repair templates were purchased from IDT. All injection reagents were diluted in water and injected at a final concentration of 50ng/μl. Worms were screened using the co-CRISPR phenotype then recombination of the repair template was screened using restriction digest and finally recombinant worms were sequenced over the targeted region to identify the mutation. The CRISPR edited line *nlg-1(qa3780)* was outcrossed against the wild-type background used in paired behavioural assays twice.

### Protein sequence alignment

Multiple protein sequence alignment of *C. elegans* NLG-1 and human NLGN1-4 was performed using the Clustal W method. *C. elegans* NLG-1 sequence was downloaded from WormBase version WS274. The longest NLG-1 isoform (C40C9.5e) was used for analysis. Human NLGN1-4 sequences were downloaded from NCBI. Accession numbers for the sequences used are as follows: NLGN1, NP_001352856; NLGN2, XP_005256801; NLGN3, NP_061850; NLGN4, AAQ88925.

## Acknowledgments

Cas9 plasmid, sgRNA’s targeting *unc-58* and *dpy-10* and *unc-58 and dpy-10* repair templates were kindly provided by Thomas Boulin. *nlg-1(ok259)* was provided by CGC, which is funded by NIH Office of Research Infrastructure Programs (P40 OD010440). This work was funded by the Gerald Kerkut Charitable Trust.

---

**Figure S1.**
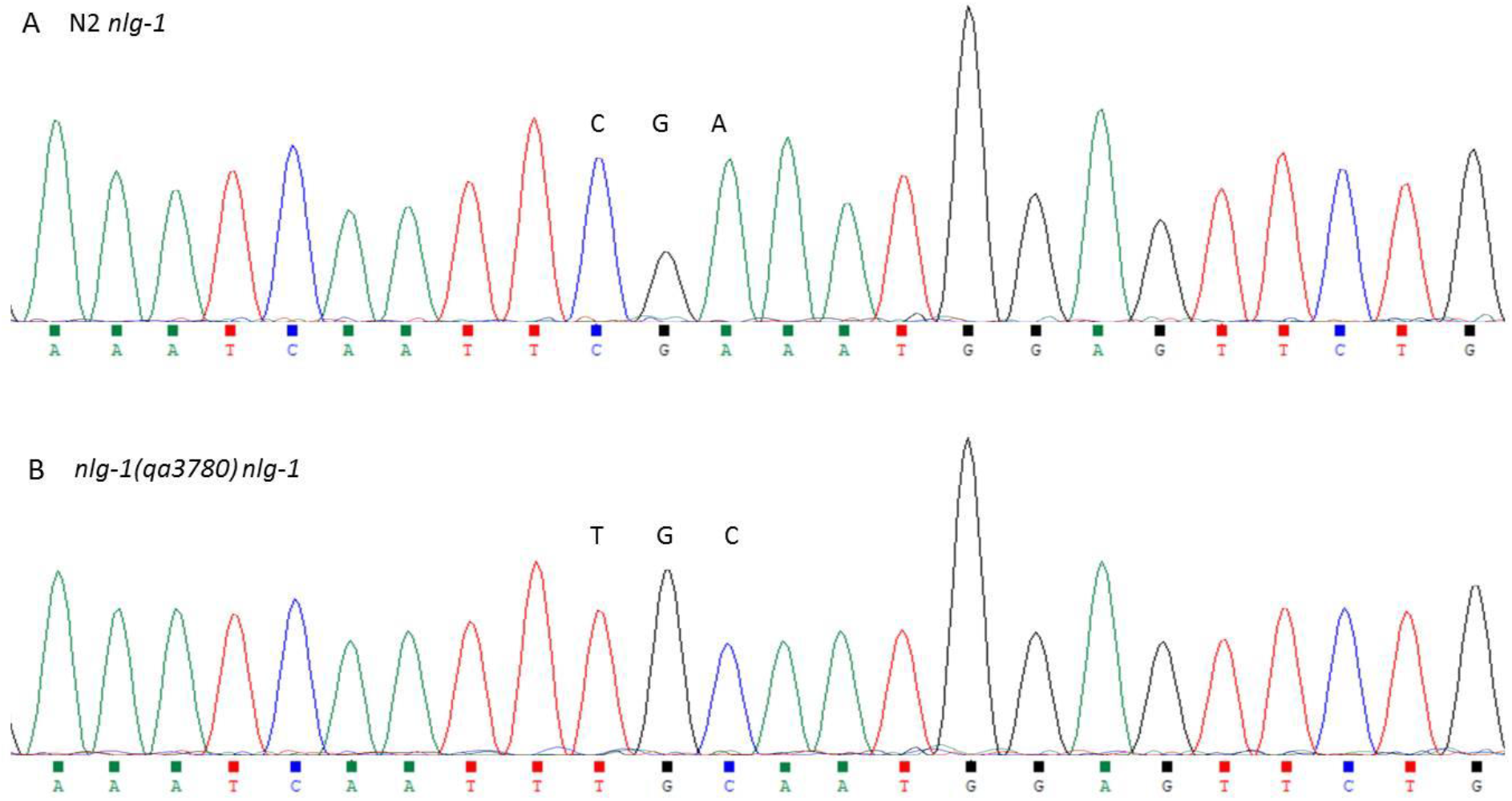
CRISPR generated *nlg-1(qa3780)* nucleotide edit. (A) Chromatogram showing partial N2 *nlg-1* DNA sequence. The codon of interest, CGA, to be edited by CRISPR is indicated. (B) Chromatogram showing partial *nlg-1(qa3780) nlg-1* DNA sequence. The edited codon, TGC, is indicated. Sequence is displayed in the 5’ to 3’ orientation.

